# Hyperspectral super-resolution imaging with far-red emitting fluorophores using a thin-film tunable filter

**DOI:** 10.1101/756023

**Authors:** Adriano Vissa, Maximiliano Giuliani, Peter K. Kim, Christopher M. Yip

## Abstract

New innovations in single-molecule localization microscopy (SMLM) have revolutionized optical imaging, enabling the characterization of biological structures and interactions with unprecedented detail and resolution. However, multi-colour or hyperspectral SMLM can be particularly challenging due to non-linear image registration issues, which affect image quality and data interpretation. Many of these arise as a consequence of differences in illumination optics (beam profile, power density, polarization, point spread function) for the different light sources. This is particularly acute in evanescent-wave based approaches (TIRF) where beam shape, decay depth, and power density are important. A potential useful approach would be to use a single excitation wavelength to perform hyperspectral localization imaging.. We report herein on the use of a variable angle tunable thin-film filter to spectrally isolate far-red emitting fluorophores. This solution was integrated into a commercial microscope platform using an open-source hardware design, enabling the rapid acquisition of SMLM images with ~ 15-20 nm spectral resolution.. By characterizing intensity distributions, average intensities, and localization frequency through a range of spectral windows, we identified an optimal fluorophore pair for two-colour SMLM. Fluorophore crosstalk between the different spectral windows was assessed by examining the effect of varying the photon output thresholds on the localization frequency and fraction of data recovered. The utility of this approach was demonstrated by hyper-spectral super-resolution imaging of the interaction between the mitochondrial protein, TOM20 and the peroxisomal protein, PMP70.

## I. INTRODUCTION

With the advent of single-molecule localization microscopy (SMLM), new limits of optical resolution have been realized, and the range of biological questions that can be addressed with fluorescence imaging has increased dramatically. Techniques such as stochastic optical reconstruction microscopy (STORM/dSTORM)^1^ and photoactivated localization microscopy (PALM)^2^ are able to provide lateral resolution of 10-20 nm with a corresponding axial resolution of 10-100 nm.^3, 4^ This enhanced resolution has facilitated the sub-diffraction limit visualization of cellular structures such as organelle ultrastructure and protein spatial distribution in both fixed and live cells, ^5–7^ Characterization of heterotypic protein-protein interactions often requires the use of multi-colour imaging strategies; however, the diverse photophysical properties of many dSTORM fluorophores^8–10^ and PALM fluorescent proteins^2, 11^ can lead to unpredictable experimental outcomes, including unequal localization precision across channels in multi-colour imaging.

Several of the far-red emitting photoswitchable fluorescent molecules commonly used in dSTORM imaging have been shown experimentally to exhibit favourable and reliable photophysical properties in a buffered reducing environment.^10, 12^ These include an inducible dark state transition, high duty cycle, moderate to high photon output, and sensitivity to 405 nm excitation.^13^ Each of these factors contributes to the blinking behaviour of the fluorophore and thus the molecular density and structural integrity of the final image. Interestingly, small molecule non-far-red emitting fluorophores often do not consistently exhibit characteristics appropriate for localization microscopy. This is problematic since conventional multi-channel imaging relies on the acquisition of data from non-overlapping spectral regions, It can therefore be challenging to collect and process multi-colour super-resolution data. Furthermore, multi-channel SMLM data is very sensitive to chromatic aberrations, which can become the limiting factor in determining the extent of colocalization uncertainty between structures imaged in two separate spectral channels. For these reasons, several groups have recently focused on developing methods for spectrally resolving overlapping far-red emission spectra.^14, 15^ In one instance, this was accomplished by using a long-pass emission filter while another relied on a prism-based technique, which enabled simultaneous collection of fluorescence intensity and spectral information. While the former offers a proven and reproducible method for resolving SMLM fluorophores based on intensity ratio, it relies on a fixed emission cutoff and requires a split-view or multiple cameras.^16^ The second approach offers true colour super-resolution and the ability to combine more than two fluorophores, but involves a non-trivial optical imaging setup. Hyperspectral imaging has also been accomplished using acousto-optic tunable filters (AOTFs) and liquid crystal tunable filters (LCTFs).^17^ Both of these particular approaches offer excellent speed and specificity, yet suffer from a significant attenuation of emission intensity, which renders them unsuitable for SMLM. A alternative approach is to use a thin-film tunable filter (TTF), which features exceptionally high transmission and very steep spectral edges.^17^ Varying the angle of incidence of emitted light on the filter enables hyperspectral imaging with ~ 15 – 20 nm spectral resolution. In this study, we examined the performance of a TTF in distinguishing between a pair of far-red SMLM fluorophores with overlapping emission spectra using a single excitation laser line and full-field hyperspectral imaging on a single EMCCD camera.

## II. EXPERIMENTAL METHODS AND CALIBRATION

### A. Optical Setup

All dSTORM imaging was performed on a home-built TIRF microscope. For this platform, a Galilean beam expander was used to expand the excitation laser (170 mW, 643 nm) to one inch in diameter. A beam steerer post was mounted on top of a linear translation stage allowing for imaging in any of three imaging modalities: epifluorescence (Epi), HiLo, or TIRF. A lens used to focus the excitation beam in the back focal plane of the objective was mounted on the steerer such that the distance between the focusing lens and the objective remained constant over the entire operational regime (Epi, HiLo, TIRF). The excitation beam entered the microscope body (dual-deck Olympus IX83, Olympus Canada, Richmond Hill Ontario) through the back port of the upper deck fitted with a standard fluorescence turret. The filter cube for these experiments only contained a 660 nm dichroic beamsplitter. For these experiments, the emission filter was removed, allowing for the full range of possible emission wavelengths. A custom optical plate holding the thin-film tunable filter (TTF: Semrock Versachrome TBP01-697/13) was placed in the lower deck of the IX-83. The TTF mount was fitted to a galvanometer, which controlled the angle of incidence (AOI) between the optical axis of the microscope (objective) and the filter normal. All images were collected on a Photometrics QuantEM EMCCD camera. Image acquisition and galvo scanning were performed with the open source software Micromanager (version 1.4.22). A schematic representation of the optical setup is presented in Figure 1(a). A basic current source was built based on a quadruple half-H driver (SN754410) that followed a TTL signal from an Arduino microcontroller (Figure 1b). The Arduino was configured to act as both the shutter control and configurable emission filter. To this end, each setting in Micromanager activated two output pins in the Arduino board. The first controlled the shutter and the other activated a channel of the half-H drive, selecting for the specific spectral region of the emission spectra.

**FIGURE 1.**
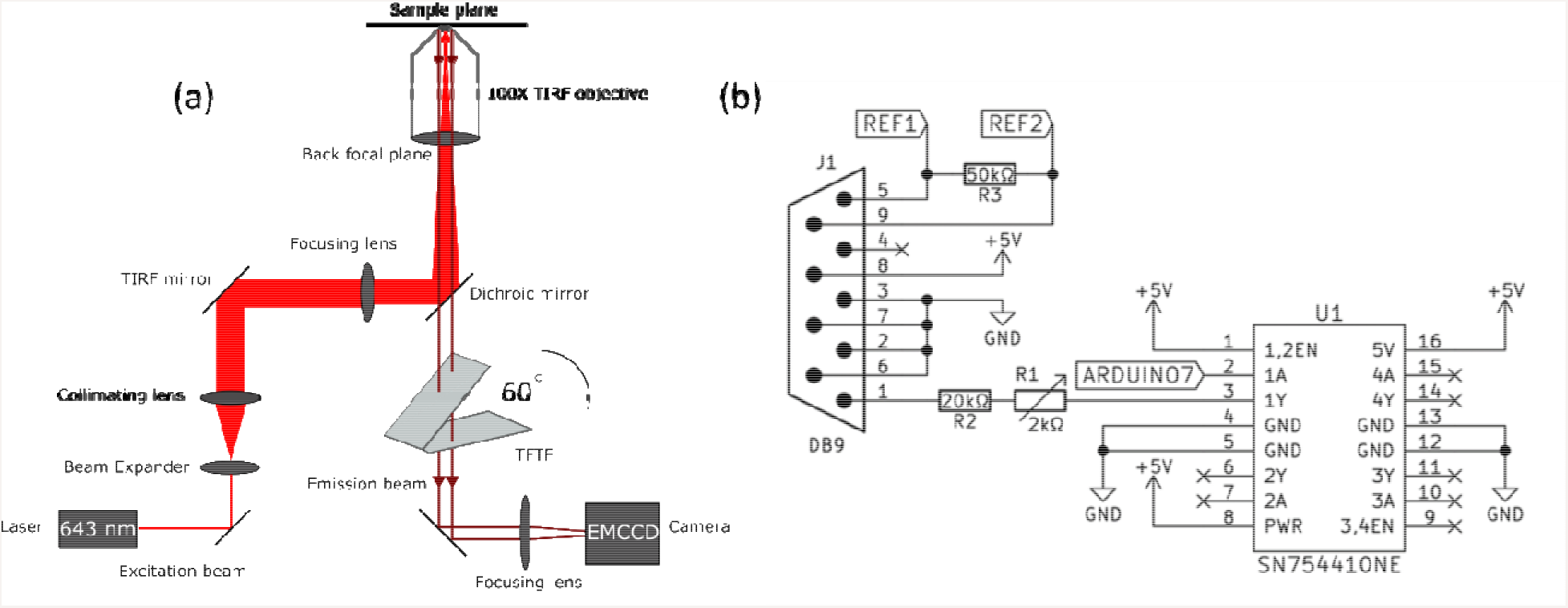
(a) Excitation light from a 643 nm DPSS laser is expanded, collimated and positioned in the back focal plane of the TIRF objective. Fluorescence emission travels back through the objective, and arrives at the TTF which has been integrated into the microscope body. Modulating the angle of incidence allows selection of the spectral window detected by the EMCCD camera. (b) The Arduino microcontroller is used to control the rotation of the galvanometer. A TTL pulse from the Arduino sends a 5V signal thought the circuit. The current can be modulated by two resistors in series (R1 and R2). The resulting voltage can be used as a reference value for angle of incidence (REF1 and REF2).

### B. TTF calibration and fluorophore characterization

Modulating the angle of incidence (AOI) from 0° to 60° allowed for the selection of ~ 20 nm wide spectral regions between 615 nm and 700 nm. To calibrate the system and ensure proper selection of the emission bands, polychromatic light was transmitted along the optical path. By varying the driving voltage to the galvanometer, and using a spectrometer (S2000, Ocean Optics, Dunedin, Florida) mounted to the side port of the microscope, a calibration curve was generated that mapped the driving voltage, AOI, and the corresponding 20 nm emission bands with center wavelengths varying from 670 nm to 700 nm. (Figure 2(a)).

**FIGURE 2.**
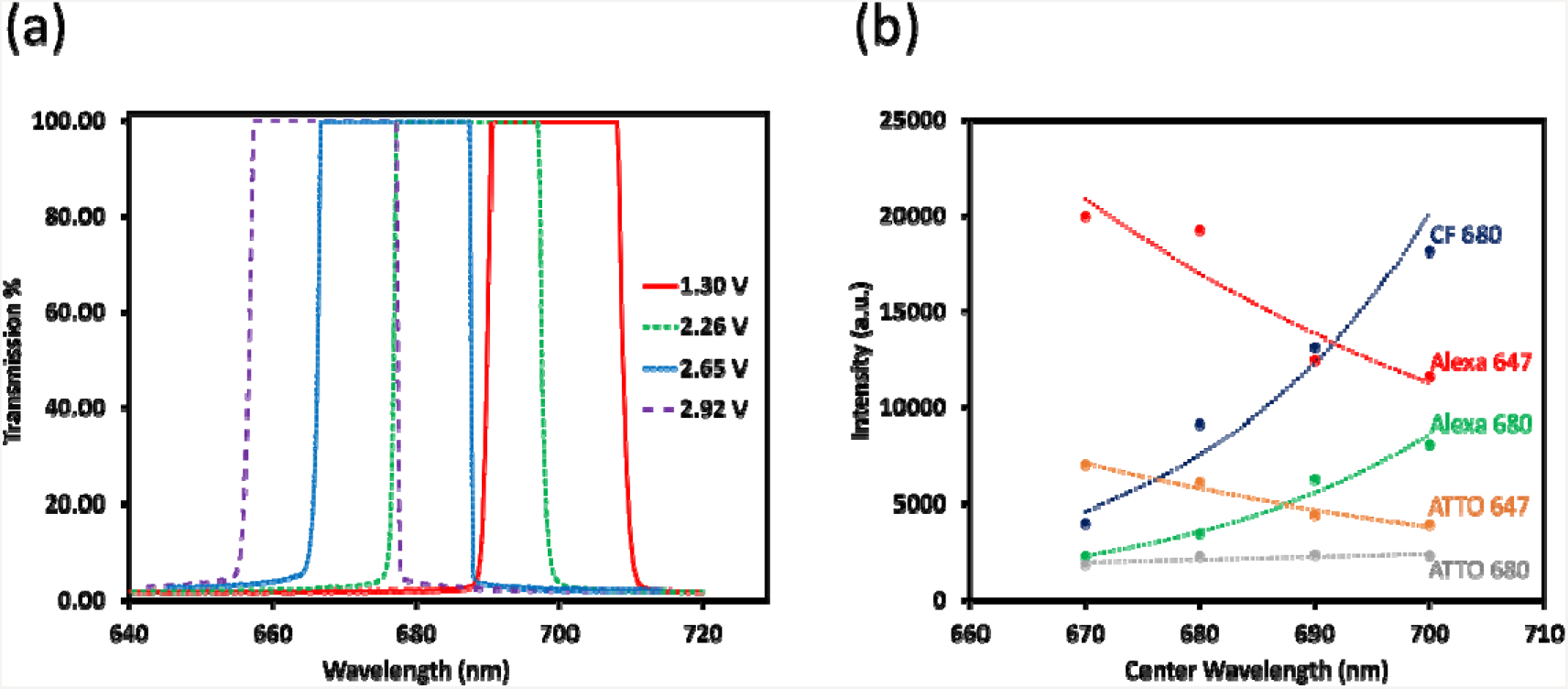
(a) TTF emission band as a function of voltage measured from the galvanometer. The emission of antibody-linked Alexa 647 was measured with a portable spectrometer at multiple output voltages to determine the transmission over a range of wavelengths. The measured voltage was then used to adjust the AOI to the desired angle and corresponding emission band. (b) Fluorescence intensity of 5 far-red emitting fluorophores across spectral windows: CW 670, CW 680, CW 690 and CW 700.

Subsequently, the emission intensity of 5 far-red emitting fluorophores (Alexa 647, ATTO 647, Alexa 680, ATTO 680 and CF 680) was measured at several points between the 0° and the 20° position of the filter by taking the average intensity of 3 TIRF images per center wavelength (Figure 2(b)). In all cases, the standard deviation of the measurements was negligible. The EMMCD gray level increased and decreased in a predictable and reproducible manner for all fluorophores, with considerable differences in the relative intensity values of the emitters across the spectrum. This initial characterization identified Alexa 647 and CF 680 as the fluorophore pair with the largest change in intensity (ΔI) measured in percentage terms as:

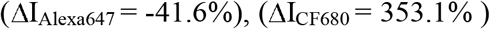

### C. Theoretical vs. experimental emission spectra

To further validate the spectral sensitivity and performance of the TTF, two colour imaging with the far-red emitting fluorophores: Alexa 647 and CF 680, was performed. Using the TTF to select spectral windows with center wavelengths of 670 nm (CW 670) and 700 nm (CW 700) respectively, isolated the theoretical emission maxima of each fluorophore (Figure 3(a)). However, in order to obtain more relevant spectral data, the emission of the same five fluorophores was measured experimentally using a side-port mounted spectrometer. The experimental spectra showed a significant deviation from the theoretical emission curves (Figure 3(b)). Consistent with intensity data shown in (Figure 2(b)), the experimental spectra revealed the variation in efficiency of each fluorophore when excited at 643 nm. Alexa 647 yielded the strongest emission in the CW 670 window, while CF 680 yielded the strongest emission in the CW 700 window. The left side of Table 1 summarizes the maximum intensity ratios of Alexa 647 : CF 680 and Alexa 647 : Alexa 680 (and vice versa) in both spectral windows. Via diffraction-limited TIRF imaging, the ratio of Alexa 647 : Alexa 680 vs Alexa 647 : CF 680 was 5.15 vs 2.12. However at CW 700, the reciprocal ratios were 0.87 vs 1.89. These measurements showed that the efficiency of Alexa 647 was such that it produced a signal of higher intensity than Alexa 680, even in the CW 700 region. This behaviour raises a potential issue for intensity-based spectral assignment. CF 680 however, yielded a stronger signal than Alexa 647 at CW 700, with a CF 680 : Alexa 647 intensity ratio of 1.89. The spectral characteristics of these fluorophores was evaluated using the TTF under dSTORM blinking conditions (Figure 3(c,d)). To this end, photoswitching of individual samples of antibody-conjugated fluorophore was induced as previously reported.^13^ Compared to diffraction-limited imaging, the changes in the intensity ratios for CW 670 (Alexa 647 : Alexa 680, Alexa 647 : CF680) increased by ~ 3 and ~2, respectively. However, at CW 700, the Alexa 680 : Alexa 647 ratio decreased to 0.50, while the CF 680 : Alexa 647 ratio decreased slightly to 1.70 (Table 1).. Taken together, these data and that shown in Figure 2(b) suggested that Alexa 647 and CF 680 represent the most promising dye pair for spectral discrimination using the TTF at CW 670 and CW 700.

**Table 1.** Fluorophore maximum intensity ratios across spectral windows.

|  | TIRF/Broadband |  | dSTORM/TTF |  |
| --- | --- | --- | --- | --- |
|  | CW 670 | CW 700 | CW 670 | CW 700 |
| AF 647 : AF 680 | 5.15 | - | 16.60 | - |
| AF 647: CF 680 | 2.12 | - | 6.28 | - |
| AF 680 : AF 647 | - | 0.87 | - | 0.50 |
| CF 680 : AF 647 | - | 1.89 | - | 1.70 |

**FIGURE 3.**
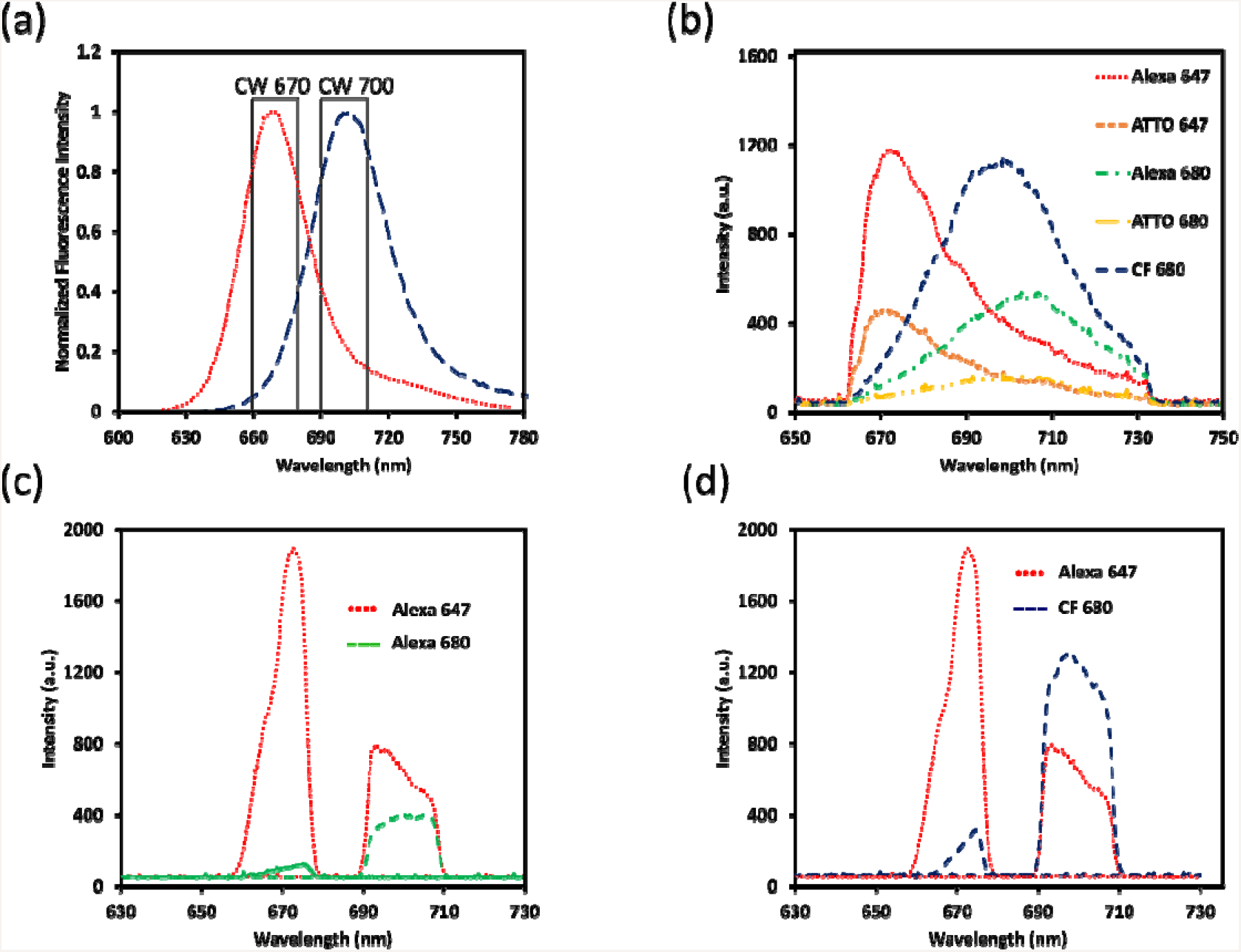
(a) Theoretical emission spectra of 647-species fluorophore (red) and a 680-species fluorophore (blue), with desired spectral windows displayed at the center wavelength of 670 nm (CW 670) and the center wavelength of 700 nm (CW 700), encompassing the emission peak maxima of the respective fluorophores. (b) Experimental emission spectra of 5 far-red emitting fluorophores collected from *in vitro* preparations of antibody-conjugated fluorophores using a spectrometer mounted to the side port of an Olympus IX-83 microscope. (c) Experimental emission spectra of Alexa 647 and Alexa 680 under photoswitching conditions at CW 670 and CW 700. Alexa 647 remains the dominant emitter in both spectral windows. (d) Experimental emission spectra of Alexa 647 and CF 680 under photoswitching conditions at CW 670 and CW 700. The emission intensity of CF 680 is larger in magnitude than that of Alexa 647 at CW 700.

### D. SMLM intensity and localization frequency across spectral windows

To characterize the intensity distributions of individual Alexa 647 and CF 680 molecules, we performed a series of dSTORM SMLM acquisitions across a range of spectral windows. The TTF angle was adjusted to collect data from 20 nm spectral bands corresponding to CW 670, CW 680, CW 690 and CW 700. Distinct variations in mean intensity and localization frequency in the histogram distribution were observed (Table 2, Figure S1). From CW 670 to CW 700, Alexa 647 displayed a mean intensity (I_avg_) decrease of 70% and localization frequency (LF) decrease of 42%. From CW 700 through to CW 670, CF 680 displayed an I_avg_ decrease of 93% and an LF decrease of 98%. The data revealed that the ΔI_avg_ and ΔLF between Alexa 647 and CF 680 was greater at CW 670 than at CW 700 (Figure 4 (a,b)), suggesting considerable Alexa 647 crosstalk at CW 700.

**Table 2.** Average intensity and localization frequency across spectral windows.

|  | CW 670 |  | CW 680 |  | CW 690 |  | CW 700 |  |
| --- | --- | --- | --- | --- | --- | --- | --- | --- |
|  | AF 647 | CF 680 | AF 647 | CF 680 | AF 647 | CF 680 | AF 647 | CF 680 |
| $I_{\text{avg}}$ (photons) | 925 | 55 | 460 | 128 | 322 | 543 | 276 | 771 |
| LF (x 1000) | 137.5 | 5.1 | 110.1 | 78.4 | 92.4 | 172.4 | 79.3 | 212.4 |

**FIGURE 4.**
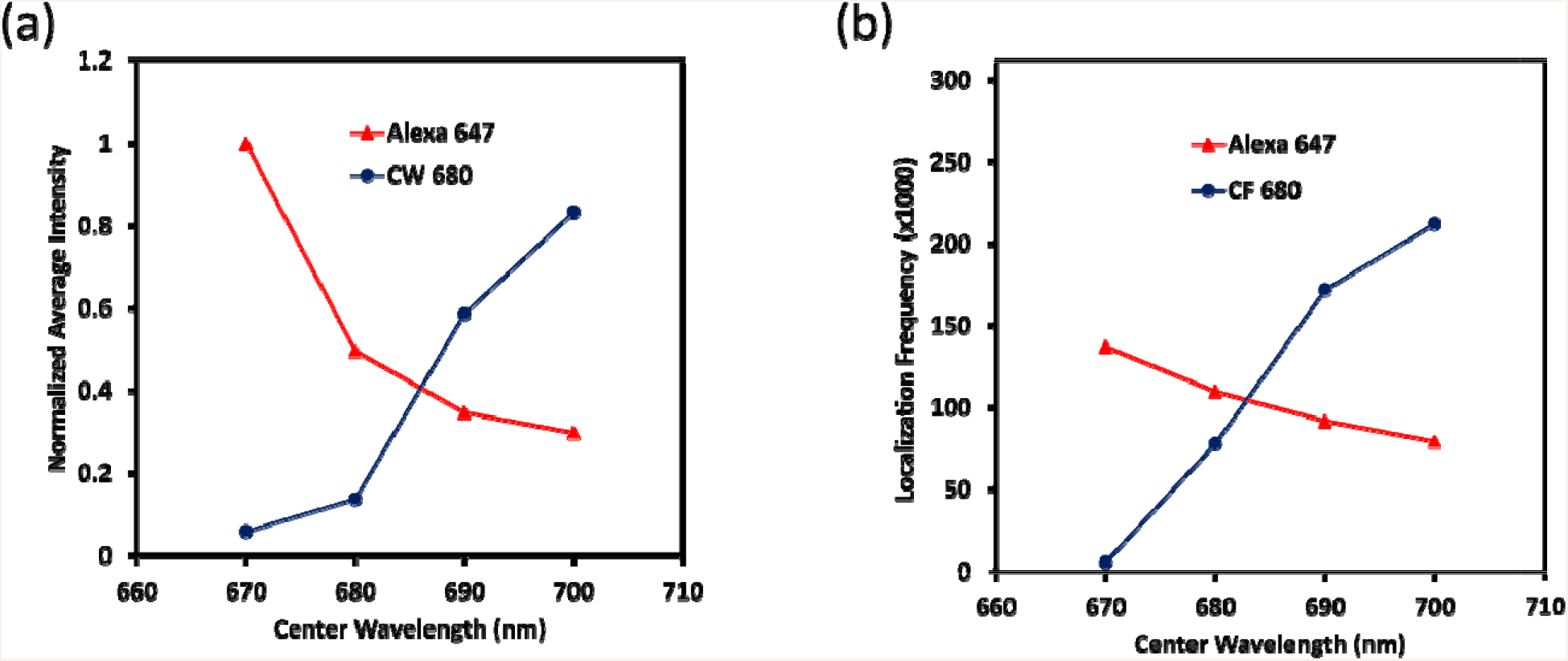
(a) Average intensity of localized molecules plotted as a function of center wavelength for Alexa 647 and CF 680. (d) Localization frequency of super-resolution reconstructions plotted as a function of center wavelength for Alexa 647 and CF 680.

## III. RESULTS

### A. SMLM imaging of control samples, thresholding and crosstalk determination

To assess how localization is affected by intensity, super-resolution images acquired for both fluorophores were analyzed at CW 670 and CW 700 over a range of photon count thresholds (Figure 5 (a,b)). As expected, a higher frequency of localizations were found for Alexa 647 at CW 670 than at CW 700. As the photon output threshold (POT) was increased from 1000 to 3000, the number of localizations decreased accordingly in both spectral windows (Figure 5 (c)). Interestingly, the opposite behaviour was observed for CF 680. The localization frequency for this fluorophore was significantly lower at CW 670 than it was at CW 700. (Table 3) However, the frequency decreased as the POT was increased, as expected. The initial frequency of localizations for CF 680 at CW 670 was very low in comparison to the other images, reflecting the extent of the attenuation of this fluorophore’s dSTORM photon output in this spectral region. The fraction of data recovered at each threshold for both fluorophores in both spectral windows was also quantified (Figure 5(d)). For a given center wavelength, the fraction recovered was calculated by dividing the localization frequency at a given threshold by the original localization frequency for the fluorophore undergoing maximal emission in that spectral window, as follows:

**Table 3.** Localization frequency (x 1000) of *in-vitro* preparations of Alexa 647 and CF 680 in spectral windows CW 670 and CW 700 at a function of photon output threshold.

| Threshold | AF 647 |  | CF 680 |  |
| --- | --- | --- | --- | --- |
|  | CW 670 | CW 700 | CW 670 | CW 700 |
| - | 137.4 | 79.3 | 5.5 | 212.4 |
| 1 | 56.3 | 4.4 | 0.17 | 60.7 |
| 2 | 16.8 | 1.9 | 0.04 | 18.0 |
| 3 | 7.4 | 0.8 | 0.02 | 8.0 |

**FIGURE 5.**
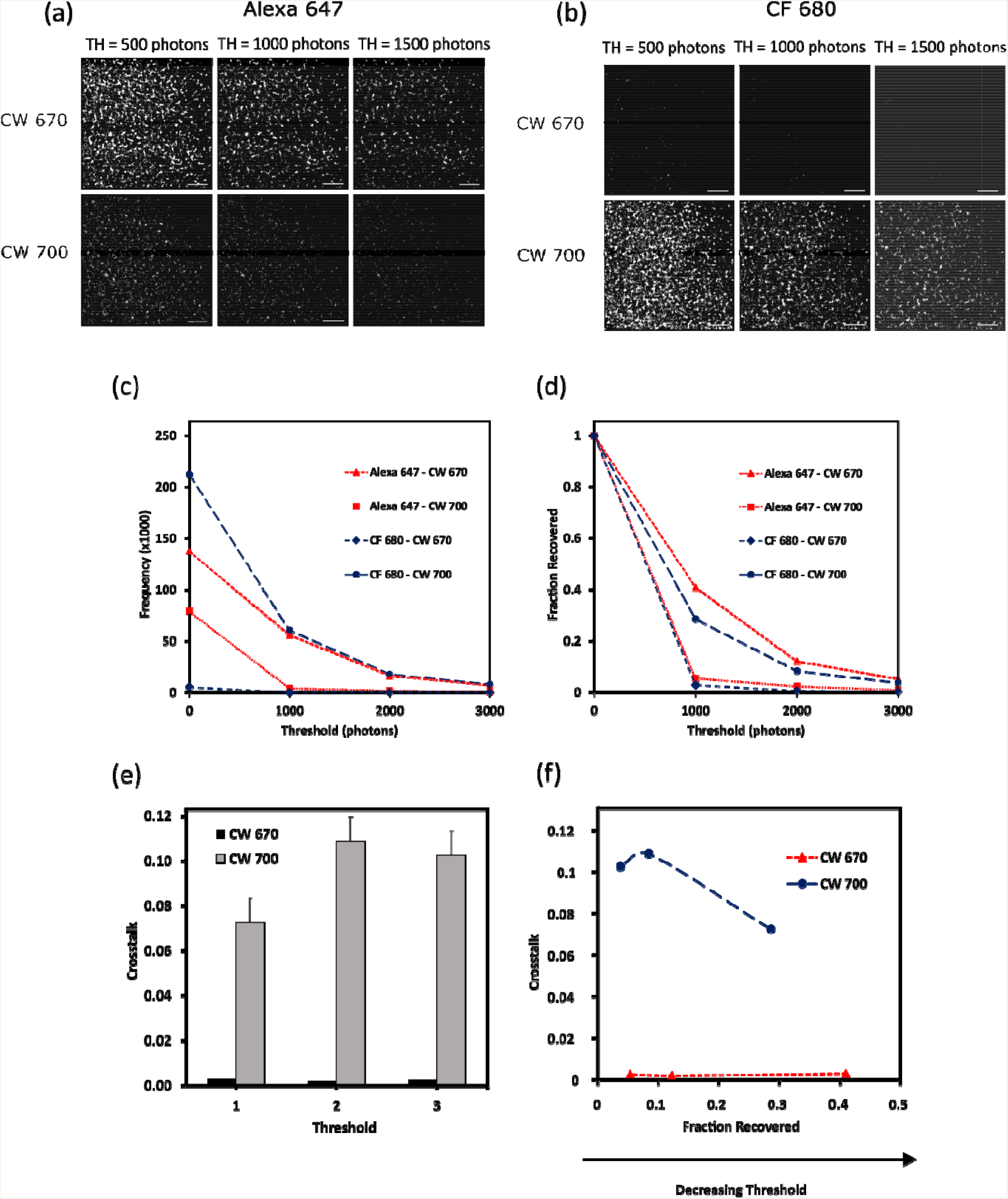
(a-b) A series of images of Alexa 647 and CF 680, respectively, at CW 670 and CW 700 displayed at various photon output thresholds (TH). Scale bars = 5 μm (c) A plot of localization frequency versus photon output threshold for all four fluorophore-spectral window combinations. (RSE = 1.93%-3.5%, N = 3) (d) A plot of fractio of localizations recovered vs. threshold for all four fluorophore-spectral window combinations (RSE = 2.2% - 3.8%, N = 3). (e) Bar graph of theoretical crosstalk at a series of thresholds for spectral windows CW 670 and CW 700. Uncertainty is expressed on each measurement as standard error,(N = 3). (f) A plot of theoretical crosstalk vs fraction of localizations recovered in spectral windows CW 670 and CW 700. (RSE = 2.3% - 3.6%, N = 3)..

For a series of thresholds *i,j,k*, and original localization frequency LF_o_

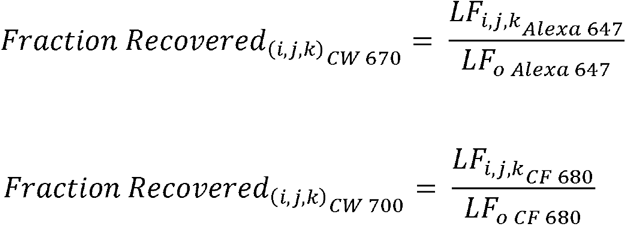

Alexa 647 and CF 680 both exhibited a similar rate of decrease in the fraction of localizations recovered as the POT was increased, as per the following:

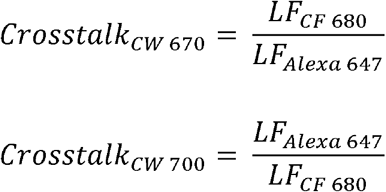

Crosstalk values at CW 670 ranged from 0.02% to 0.03% at the three photon output thresholds, again reflecting the small number of CF 680 localizations that were recovered. At CW 700, crosstalk values ranged from 7% to 11% (Figure 5(e)). Plotting crosstalk vs. fraction recovered for each spectral window at a series of thresholds provided a measure of comparison between two opposing goals: Maximizing the localization frequency of the desired fluorophore while minimizing crosstalk (Figure 5 (f)). Interestingly, at CW 700, a higher recovered fraction of CF 680 (i.e. a lower photon output threshold) was largely associated with a lower crosstalk value. While this may appear counter-intuitive, it reflects the fact that, as the POT is increased, the ratio of localized Alexa 647 molecules to localized CF 680 molecules approaches 1:1. Since Alexa 647 is a more efficient photon emitter, its contribution to crosstalk increases with increasing POT.

### B. 2-channel hyperspectral SMLM imaging of mitochondria and peroxisomes

To demonstrate the utility of the approach described herein, hyperspectral super-resolution imaging of the mitochondrial protein TOM20, labeled with Alexa 647 (TOM-Alexa 647) and the peroxisomal protein, PMP70, labeled with CF-680 (PMP70-CF 680) in HeLa cells. Images were reconstructed at CW 670 and CW 700. At CW 670, there was virtually no visible crosstalk, as seen in the *in-vitro* control and single-labeled samples (Figure S2). At CW 700, significant crosstalk was apparent. The difference in morphology between peroxisomal puncta (yellow arrows) and residual mitochondrial signal (purple boxes) served to illustrate the extent of this phenomenon (Figure 6(a)). When a POT of 300 was applied, close examination of the CW 700 image revealed that crosstalk was not completely eliminated. Increasing the threshold further may attenuate crosstalk to a greater degree, but at the expense of the true CF 680 signal. Continuously increasing the threshold with the goal of attenuating all crosstalk presents a dilemma wherein the reduction of false positives to zero is achieved only by increasing false negatives to an unacceptable level. The result is an image that lacks adequate structural context. In general, after thresholding, the Alexa 647 signal at CW 700 was comprised of sparsely situated localizations that could be only be accounted for with a density filter Figure 6(b)). The parameters of the filter used in this case were a minimum # neighbors = 5, radius = 100 nm. Under these conditions, all localizations which did not have at least 5 adjacent localizations within a 100 nm radius were removed. This served to eliminate the majority of crosstalk in the image. The CW 670 featuring TOM20-Alexa 647 and CW 700 PMP70-CF 680 images were then merged to create a two-colour hyperspectral SMLM image (Figure 6(c)).

**FIGURE 6.**
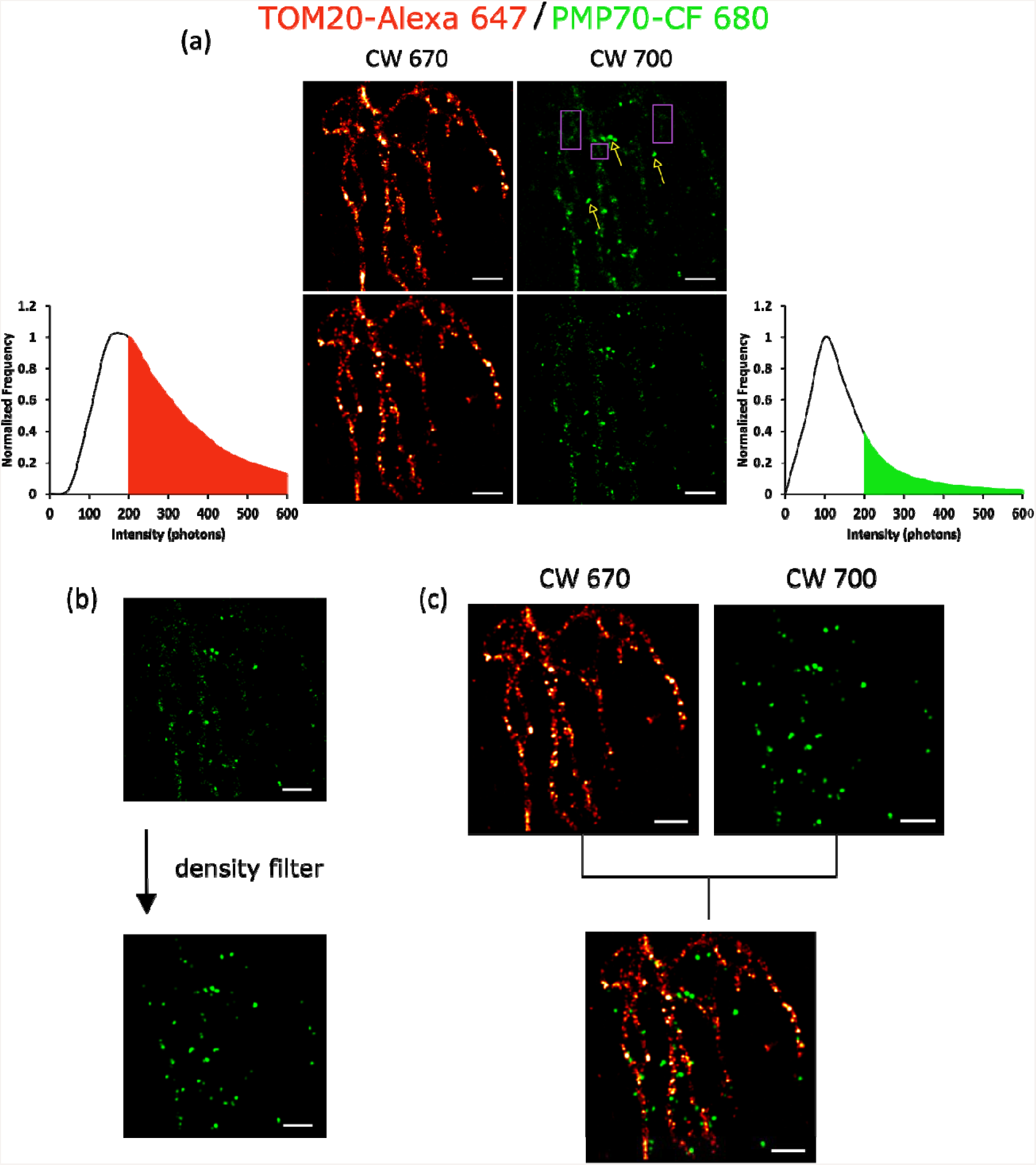
(a) Images of TOM20-Alexa 647 / PMP70-CF 680 at CW 670 and CW 700. Representative peroxisomal puncta in CW 700 are denoted with yellow arrows. Representative regions of crosstalk in CW 700 are denoted in purple boxes. A POT threshold of 200 is used to select for the desired signal while preserving the ultrastructure of the organelle. (b) Residual crosstalk that remains at CW 700 after thresholding may be removed by applying a density filter to remove sparse localizations. (c) A two-colour merged SMLM image of mitochondria and peroxisomes derived from the individual CW 670 and CW 700 images. Scale bars = 2 μm.

## IV. CONCLUSION

We have demonstrated the usefulness of a tilt-tunable filter for performing hyperspectral SMLM. This approach allowed for dSTORM SMLM of Alexa 647 and CF 680 using a single excitation source. Photon output thresholding and density filtering were used to reduce spectral cross-talk between these two fluorophores. Building from this platform, there are a number of directions to be pursued. Since biological structures are inherently heterogeneous, the absolute concentration of fluorophores in any given field of view cannot be easily determined. Therefore, although similar results were observed for multiple replicates of the data presented, a more robust statistical treatment of the data may be required to standardize photon output thresholds for specific fluorophore-protein combinations. In addition, fluorophore attribution would be facilitated by the presence of paired localizations in the two spectral windows. Simultaneous acquisition of both signals in distinct spectral windows (for example with a fixed beamsplitter and 2 EMCCD cameras or a split-view configuration) would allow for ratiometric determination of fluorophore identity.^14^ This would require the TTF to transition between CW 670 and CW 700 at a rate double that of the single localization with the shortest duty cycle in any given frame. Further experiments with optimized galvanometer stability are required to explore this possibility. In addition, modulating both the position of the spectral window in the emission region as well as its bandwidth, by using a combination of long-pass and short-pass tunable filters offers the possibility of achieving enhanced better spectral resolution between two fluorophores.

## SUPPLEMENTARY FIGURES

**FIGURE S1.**
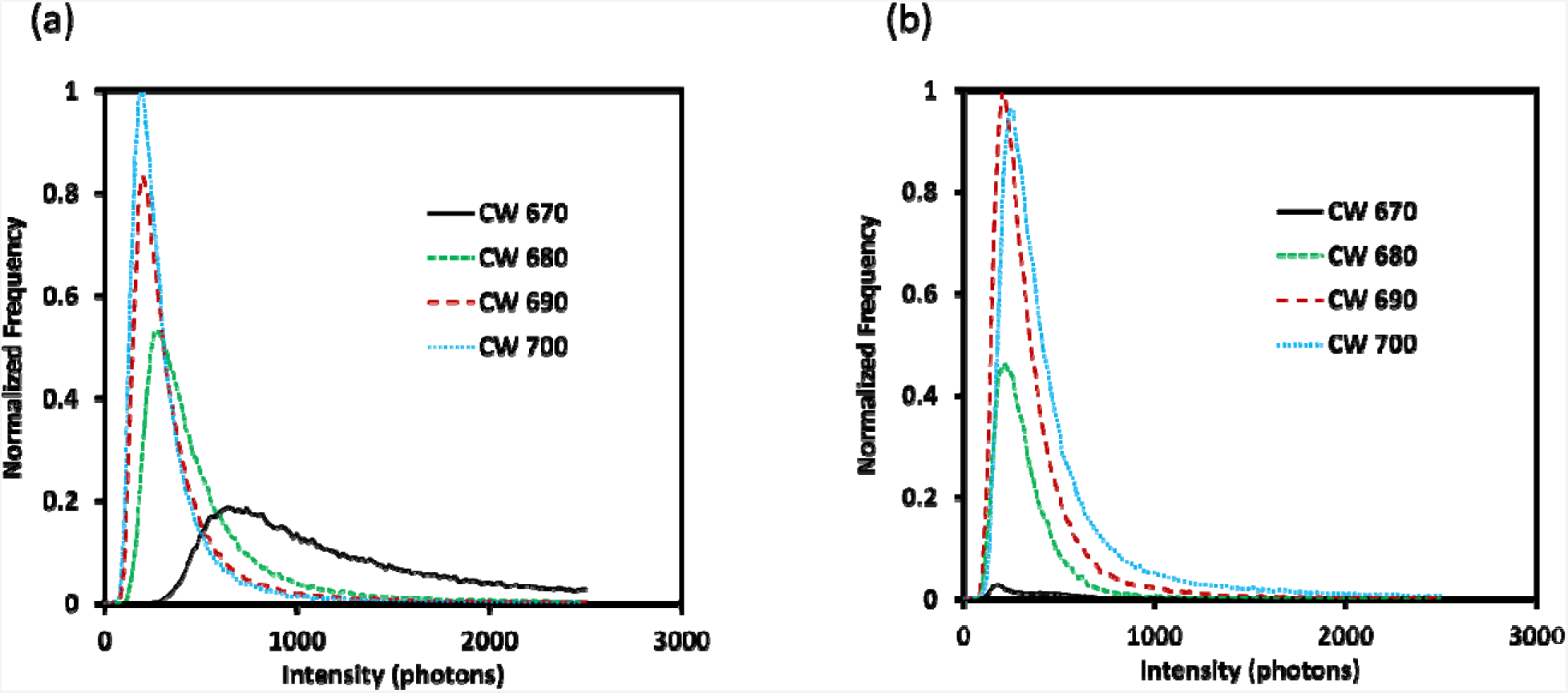
(a) SMLM intensity histograms of Alexa 647 at four spectral windows spanning the range from CW 670 to CW 700. (b) SMLM intensity histograms of CF 680 at four spectral windows spanning the range from CW 670 to CW 700. The ability to spectrally distinguish these fluorophores from each other in a mixed sample at a given center wavelength would depends on both on the mean and standard deviation of the respective intensity histograms (the distribution of data along the x-axis) and the number of detected molecules exceeding a threshold (affecting primarily the height of the histogram). A clear shift in intensity distribution provides a photon cutoff value on which to base the selection of localizations in each spectral window.

**FIGURE S2.**
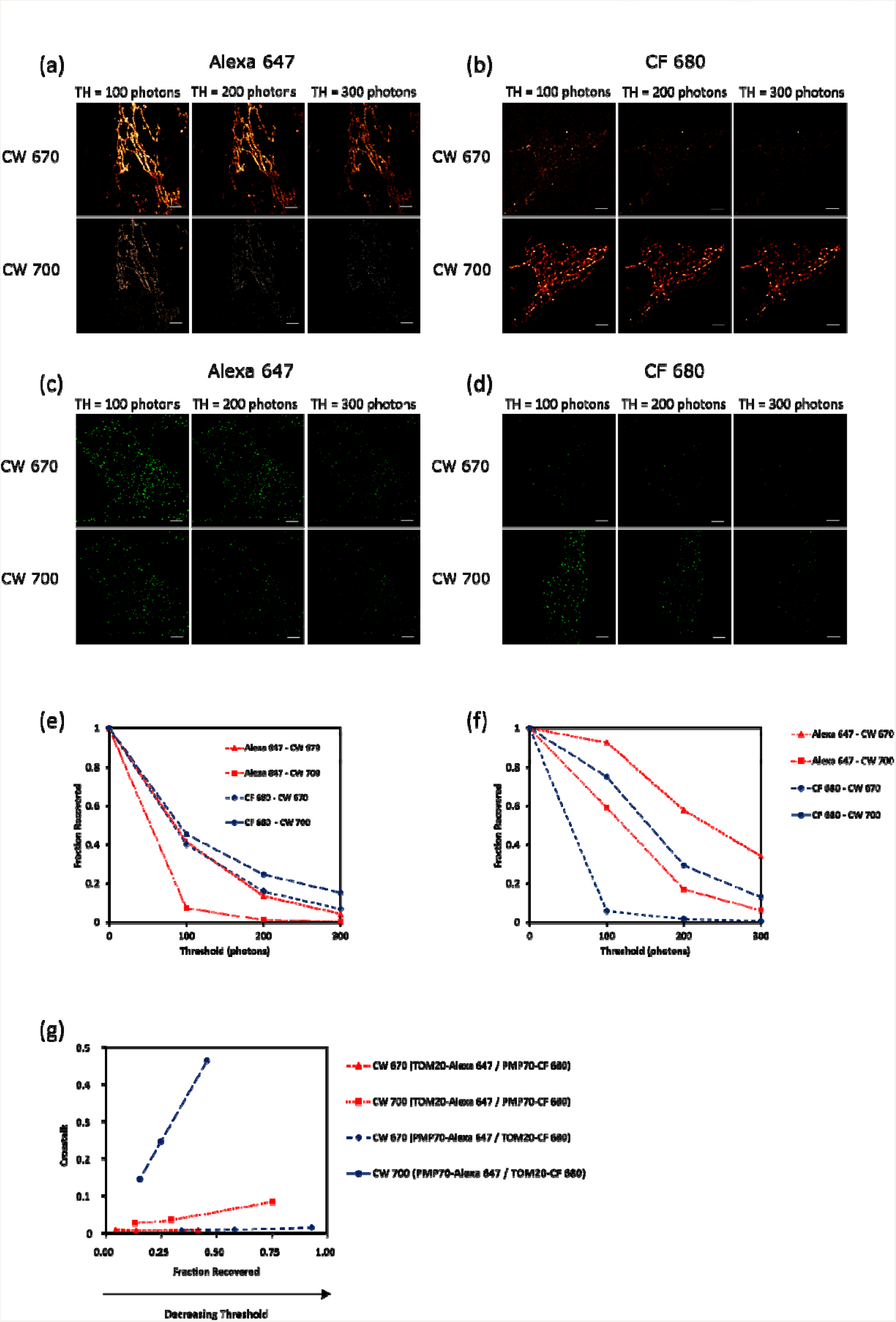
(a,b) A series of images of TOM20-Alexa 647 and TOM20-CF 680 at CW 670 and CW 700 displayed at various photon output thresholds. (c,d) A series of images of PMP70-Alexa 647 and PMP70-CF 680 at CW 670 and CW 700 displayed at various photon output thresholds. (e) A plot of fraction of localizations recovered vs. threshold for all four fluorophore-spectral window combinations in these TOM20-labeled samples. (f) A plot of fraction of localizations recovered vs. threshold for all four fluorophore-spectral window combinations in these PMP70-labeled samples. A plot of hypothetical crosstalk vs fraction of localizations recovered in spectral windows CW 670 and CW 700. The data shows that crosstalk is of greater magnitude in general in CW 700 and in particular when Alexa 647 is used to label PMP70. Scale bars = 3 μm.

